# Semaglutide interferes with postnatal development of juvenile mice with compromised growth

**DOI:** 10.1101/2024.05.06.592704

**Authors:** Amélie Joly, Lucas Rebiffé, Yves Dusabyinema, Julien Dellinger, Karine Gauthier, François Leulier, Filipe De Vadder

## Abstract

GLP-1 receptor agonists, including semaglutide, are increasingly used to treat obesity—and to a lesser extent, type 2 diabetes—in children and adolescents. Despite their therapeutic promise, the impact of semaglutide on key endocrine and metabolic developmental processes during this period is not yet fully understood. To address the limited understanding of GLP-1 receptor agonists during development, we investigated the effects of chronic semaglutide treatment on growth and metabolism of weanling male mice. In juvenile mice displaying a normal growth pattern, semaglutide lowered blood glucose without affecting food intake or weight gain. In contrast, chronic semaglutide treatment in growth-stunted juveniles exacerbated linear growth impairment, despite preserving glucose-lowering effects. Transcriptomic analysis of the liver, complemented by gene expression studies in other metabolically active tissues, suggested altered energy expenditure, indicating a complex interaction between semaglutide, energy metabolism, and growth during juvenile development. These findings underscore the importance of developmental context when evaluating the safety and efficacy of GLP-1 receptor agonists in pediatric populations.

## Introduction

In recent years, GLP-1 receptor agonists (GLP-1RAs) have emerged as significant therapeutic options for managing type 2 diabetes and obesity. As of 2025, seven GLP-1RAs are approved by the US Food and Drug Administration for the treatment of type 2 diabetes^1^. Among them, semaglutide was optimized to have a longer half-life in humans and a better resistance to Dipeptidyl Peptidase IV inhibitors than other GLP-1RAs^2^. Semaglutide improves glycemic control through pleiotropic mechanisms, including increased insulin secretion in response to glucose stimulation and improved beta cell survival and proliferation^3^. Besides its effects on glucose homeostasis, semaglutide significantly lowers energy intake and induces remarkable weight loss^1,4^. Hence, it is now widely used to treat obesity in adults^5,6^, with growing evidence supporting its efficacy in children as well^7,8^. Although semaglutide has been widely studied in adult mouse models of diet-induced obesity, its effects during early developmental windows, such as infancy and adolescence, remain poorly understood. Yet, treatment timing may be critical, as semaglutide profoundly reshapes energy homeostasis.

The juvenile period, a phase characterized by intensive growth both in humans and mice, is a highly energy-demanding process. In mammals, linear growth is regulated by the somatotropic axis, governed by growth hormone (GH) and insulin-like growth factor 1 (IGF-1). Interestingly, both *in vitro* and *in vivo* work in rodents demonstrated that GLP-1RAs can promote growth-related phenotypes by enhancing osteoblast proliferation and activity^9,10^. Besides, several studies in mice have shown that GLP-1 and GLP-1RA may enhance IGF-1 receptor (IGF-1R) signaling pathways^11,12^, favor bone metabolism homeostasis^9,13^ and even directly cross-activate the IGF-1R in human cardiac fibroblasts^14^. In addition, short- or long-term administration of GLP-1RAs in healthy adult volunteers has been shown to increase circulating GH levels^15^. Thus, it has been suggested that GLP-1RAs, such as semaglutide, could interact with the somatotropic growth axis^16,17^.

Despite these insights, how GLP-1RAs affect energy balance and growth at steady state during the juvenile period remains largely unexplored. In this study, we examined the effects of chronic semaglutide treatment on growth and energy metabolism in non-diabetic juvenile mice under optimal and impaired growth conditions. We found that semaglutide improves glucose clearance of non-obese, non-diabetic juvenile mice, irrespective of their growth phenotype. However, while it does not affect growth in normally developing mice, semaglutide further compromises linear growth in stunted juveniles. Notably, these growth-inhibitory effects appear largely independent of alterations in the somatotropic axis. Transcriptomic profiling of the liver, completed by gene expression analyses in other metabolically active organs, suggests that semaglutide may impair growth in stunted juveniles by downregulating pathways involved in basal energy expenditure and anabolic metabolism.

## Materials and methods

### Animals and treatments

All animal procedures have been approved by a local animal care and use committee (CECCAPP) and subsequently authorized by the French Ministry of Research (APAFIS #40481-2022103017295302 v3).

21-day-old C57BL/6N male mice were purchased from Charles River and weaned onto either the standard AIN-93G diet (Control Diet, CD) or on an isocaloric, protein-restricted modification of the AIN-93G designed to induce stunting (Stunting-Inducing Diet, SID), as previously described in our study^18^. The detailed composition of both diets is provided in Table 1. Animals were weighted and body size was measured weekly under brief isoflurane anesthesia. They received subcutaneous injections of semaglutide (Ozempic®, Novo Nordisk 0.5 mg injectable solution) at a dose of 30 nmol/kg body weight, or vehicle (0.9% NaCl), twice weekly until postnatal day 56 (P56). Injections were administered one day prior to the oral glucose tolerance test (OGTT), and one day before sacrifice.

At P56, animals were fasted for 6 hours starting at 8 AM and killed by cervical dislocation at 2 PM. Blood and organs were sampled. The left femur and tibia were dissected and measured precisely using a caliper.

For OGTT, mice were fasted for 6 hours starting at 8 AM and gavaged with a 20% D-(+)-glucose solution (Sigma Aldrich, G7021) at 1 g/kg body weight. Blood glucose was measured from the tail vein at 0 (baseline), 15, 30, 60, 90, and 120 min post-gavage using a handheld glucometer (OneTouch Verio Reflect, LifeScan Europe). Blood samples were collected from the tail vein at 0, 15 and 30 min, serum was extracted and stored at –20°C until further processing.

Only male mice were included in this study, as we previously demonstrated that SID induces greater growth impairment in juvenile males compared to females^18^.

### Insulin measurements

Serum insulin levels were measured using the Ultra Sensitive Insulin ELISA kit (Crystal Chem, 90080), following the manufacturer’s instructions.

### Western blotting

Frozen liver samples were processed for Western blotting as described by Schwarzer et al.^19^. Briefly, samples were lysed in RIPA Plus lysis buffer (50 mM Tris, 1% IGEPAL, 0.5% sodium deoxycholate, 0.1% SDS, 150 mM NaCl, 2 mM EDTA, 50 mM NaF, protease inhibitors). Protein concentration was measured using the BCA assay (Thermo Scientific, 23227). Equal amounts of protein (30 µg per well) were separated in Mini-PROTEAN TGX Stain Free precast gels (Bio-Rad, 4868096), and transferred to nitrocellulose membranes using the Trans-Blot Turbo system (Bio-Rad). Membranes were blocked and incubated for 1 hour at room temperature with either rabbit monoclonal anti-total Akt (C67E7) (1:1,000, Cell Signaling, #4691), or rabbit monoclonal anti-phospho S473 Akt (D9E) (1:2,000, Cell signaling, #4060). After washing, membranes were incubated for 1 hour at room temperature with HRP-conjugated anti-Rabbit IgG secondary antibody (1:2,500, Promega, W4011). Detection was performed using ECL Western Blotting Substrate (Promega, W1001), and signals were captured with the Amersham IQ800 imager (Cytiva). Quantification was performed using Image Lab software version 6.1 (Bio-Rad), using total protein normalization^20^.

### IGF-1 quantification

Liver tissues were weighed and homogenized in lysis buffer containing PBS, 1% Triton X100 and protease inhibitors (Roche, cOmplete™, EDTA-free). Homogenates were centrifuged (12,000 x g, 10 min, 4°C), supernatants were used to quantify IGF-1 levels using the Mouse/Rat IGF-1 Quantikine ELISA (R&D systems, MG100). Total protein content was measured using the BCA assay for normalization. IGF-1 concentrations are expressed as pg/μg total protein. The same kit was used to quantify circulating IGF-1.

### Gene expression analysis

Total RNA was extracted from brown adipose tissue (BAT) and subcutaneous white adipose tissue (scWAT) using TRI Reagent (Invitrogen, M9738). Briefly, tissues were homogenized in TRI reagent, centrifuged to remove debris, and RNA was isolated via chloroform phase separation followed by isopropanol precipitation. For liver and skeletal muscle, total RNA was extracted using the Nucleospin RNA kit (Macherey-Nagel, 740955.250).

Reverse transcription was performed on 1 μg of RNA using the SensiFAST cDNA Synthesis Kit (Meridian Biosciences, BIO-65053). Quantitative PCR was performed using Takyon No ROX SYBR Mastermix blue dTTP (Eurogentec, UF-NSMT-B0701) using a Bio-Rad CFX96 apparatus. Analysis followed MIQE guidelines^21^. Amplification efficiency was evaluated for each primer pair using standard curve assays performed prior to experimental runs. All reactions were carried out in technical duplicates to ensure reproducibility. At least two validated reference genes were used for normalization, with gene stability confirmed using the geNorm algorithm available within the CFX96 software. Primers sequences for all reference and target genes are listed in Table 2. Data are expressed as geometric means of relative expression normalized to the reference genes.

### Liver RNA-seq analysis

The concentration and quality of 10 DNAse-treated RNA samples (from SID-fed juveniles; 5 vehicle-treated and 5 semaglutide-treated) were assessed using the Qubit 4.0 fluorometer (Thermo Fisher Scientific) and the Tapestation 4150 system (Agilent Technologies), respectively. For each sample, mRNA was purified from 1 µg of total RNA using oligo(dT)-based magnetic bead isolation (Poly(A) mRNA Magnetic Isolation Module, New England Biolabs), following the manufacturer’s protocol.

Single-indexed libraries were built using the CORALL Total RNA-Seq Library Prep Kit (Lexogen), according to the manufacturer’s instructions. This fragmentation-free protocol ensures full-length transcript coverage. Unique Molecular Identifiers (UMIs; 12bp tags) were introduced during the linker Ligation step, enabling the identification and removal of PCR duplicates during downstream analysis to reduce amplification bias.

After quality control (Qubit 4.0 and Tapestation) analysis, libraries were pooled together in equimolar concentrations and sequenced on two high-output runs in a single-end mode (1×86 bp) using a NextSeq500 Illumina sequencer (Illumina). A total of 1.082 billion reads were generated, yielding an average of 36 million reads per sample (range: 31-44 million), after demultiplexing.

Read preprocessing was performed using a custom pipeline derived from Lexogen’s recommendations. Briefly, adapter trimming was performed using the Cutadapt package (version 3.5)^22^ and read quality was assessed using FastQC. Read mapping was achieved with STAR (Spliced Transcripts Alignment to a Reference, version 2.7.11a)^23^ using the *Mus musculus* GRCm39 genome reference (Ensembl, release 110). UMI-based deduplication was carried out using UMI-tools dedup (version 1.1.2)^24^, enabling accurate removal of PCR duplicates. Gene-level quantification was performed with htseq-count (version 2.0.3)^25^, generating raw count tables for each sample.

Differential analysis was performed in R (v.4.3.1), using the DESeq2 package (v.1.42.0)^26^. Visualization of differential gene expression was done using the Enhanced Volcano package (v.1.20.0)^27^, applying cut-offs of adjusted p-value (FDR) ≤ 0.01 and |log_2_ Fold Change| ≥ 1. One vehicle-treated sample was excluded from analysis following unsupervised clustering and quality control, as it appeared to be an outlier.

The RNA-seq dataset has been deposited in the NCBI Gene Expression Omnibus^28^ (GEO) under accession number GSE266477 (https://www.ncbi.nlm.nih.gov/geo/query/acc.cgi?acc=GSE266477).

### Statistical analysis

Statistical analysis was performed using GraphPad Prism v10.2.1 and R v. 4.4.3. For comparisons between two groups, unpaired *t*-tests were used when data met assumptions of normality and homogeneity of variance; otherwise, non-parametric Mann-Whitney *U*-tests were applied.

Longitudinal data—including body weight, linear size, and glycemia during the OGTT—were analyzed using linear mixed-effects models in R, employing the lme4 (v. 1.1-4), lmerTest (v. 1.1-36), and emmeans (v. 1.11.0) packages. Mouse ID was included as a random effect to account for repeated measures. Two independent experimental batches were used, and batch was included in all models as a fixed effect along with its interactions with treatment and age. Although the fixed effects involving batch were not statistically significant (p > 0.05), model comparison showed that including the full Treatment × Age × Batch interaction significantly improved model fit. Additionally, the variance attributed to batch as a random effect was consistently small. Therefore, to account for possible batch-related structure while minimizing visual redundancy, representative data from one batch are shown in the figures. Linear mixed-effects model results are reported in Table 3.

## Results

### Chronic semaglutide treatment improves glucose tolerance without altering growth of non-diabetic non-obese juvenile mice

To determine the effects of chronic semaglutide treatment during the juvenile developmental window, we characterized glucose metabolism of animals fed an experimental diet meeting the nutritional requirements for optimal growth conditions (AIN-93G^29^, hereafter referred to as control diet, CD), and treated with either vehicle (Veh) or semaglutide (Sema). Consistent with findings in adult mouse models of obesity^30^, chronic semaglutide treatment lowered fasting glycemia and insulinemia in juvenile mice (Fig. 1A-B). Semaglutide also enhanced glucose clearance during an OGTT (Fig. 1C), likely reflecting increased insulin sensitivity (Fig. 1D). In contrast to its known effects in mouse models of diet-induced obesity^31^, chronic semaglutide treatment did not reduce body weight or food intake in juvenile animals maintained under optimal nutritional conditions (Fig. 1E-F). Linear growth was also unaffected, as shown by similar body size (Fig. 1G), and bone measurements (femur and tibia lengths; Fig. 1H-I) at the end of the treatment. Assessment of the somatotropic axis revealed no detectable changes in circulating GH/IGF-1 signaling pathway (Fig. S1), consistent with the absence of growth alterations. Furthermore, semaglutide treatment did not impact body composition, with similar weights of subcutaneous white adipose tissue and gastrocnemius muscle between groups (Fig. 1J-K). Altogether, these results show that chronic semaglutide treatment modulates glucose metabolism in healthy juvenile mice without affecting growth dynamics or body composition.

**Figure 1.**
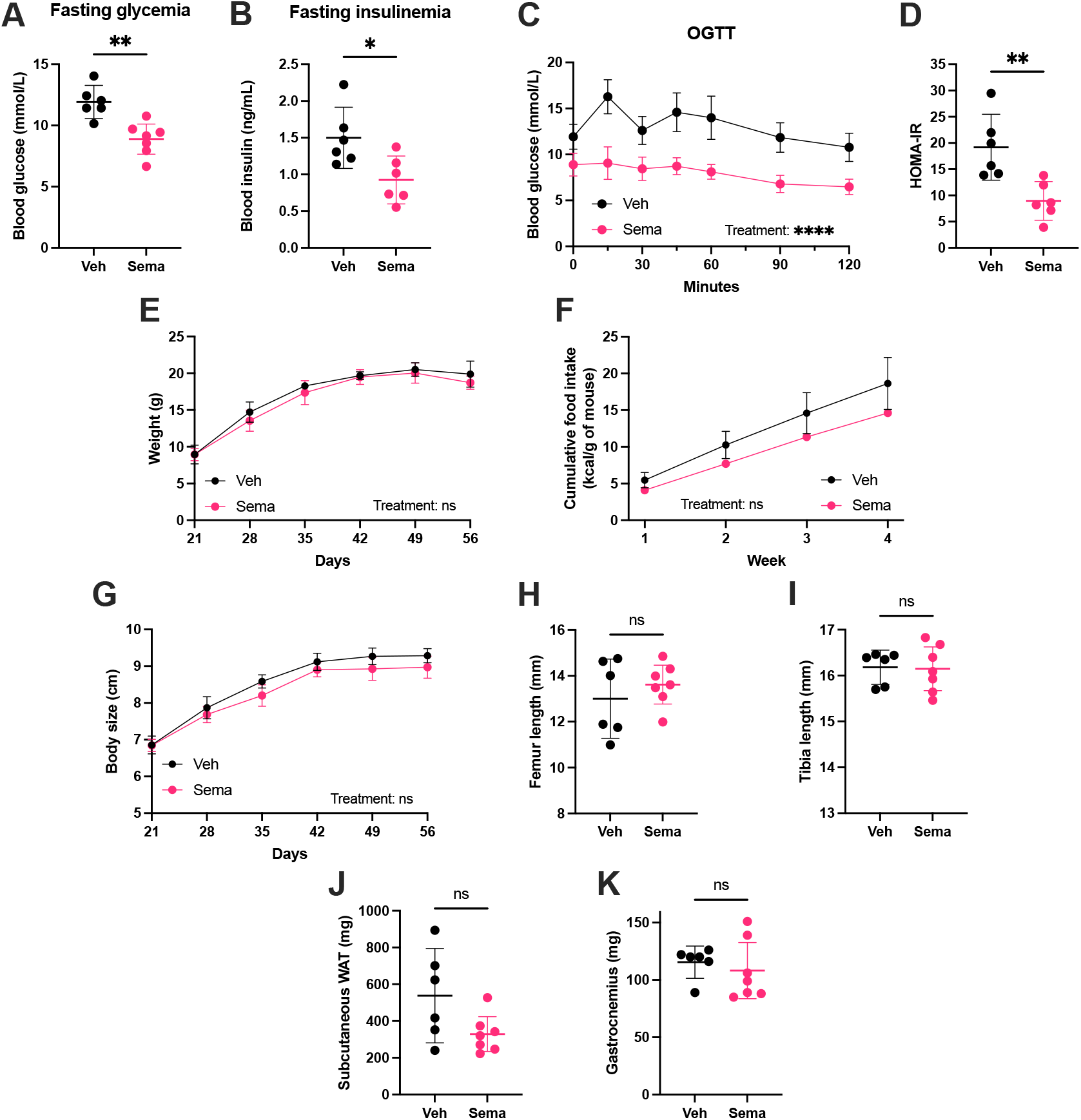
Semaglutide modulates glucose metabolism without affecting growth in control-fed juvenile mice. (A) Fasting glycemia and (B) insulinemia. (C) Glycemia during an oral glucose tolerance test (OGTT). (D) HOMA-IR. (E) Body weight over time. (F) Cumulative food intake. (G) Body size. (H-I) Femur (H) and tibia (I) lengths at postnatal day 56 (P56). (J-K) Weights of subcutaneous white adipose tissue (WAT) (J) and gastrocnemius muscle (K) at P56. Veh, vehicle-treated; Sema, semaglutide-treated. Data are shown as mean ± 95% CI, except for cumulative food intake (mean ± SEM). *p < 0.05, **p < 0.01, ****p < 0.0001, ns: not significant.

### Chronic semaglutide treatment exacerbates growth impairment in stunted juvenile mice

Given the importance of linear growth as a hallmark of juvenile physiology and the observed metabolic modulation by semaglutide under optimal nutritional conditions, we next investigated its effects in a context of disrupted growth. Stunting, defined as low height-for-age, was induced in juvenile males by feeding a low-protein variant of the AIN-93G (referred to as the stunting-inducing diet, SID; Table 1)^18^. Unexpectedly, semaglutide treatment further impaired linear growth in SID-fed juveniles, resulting in a 7% reduction in body size compared to vehicle-treated controls, along with corresponding reductions in femur and tibia length (Fig. 2A-C). These findings indicate that semaglutide specifically disrupts body development under a growth-limiting nutritional environment, highlighting the context-dependency of its physiological effects.

**Figure 2.**
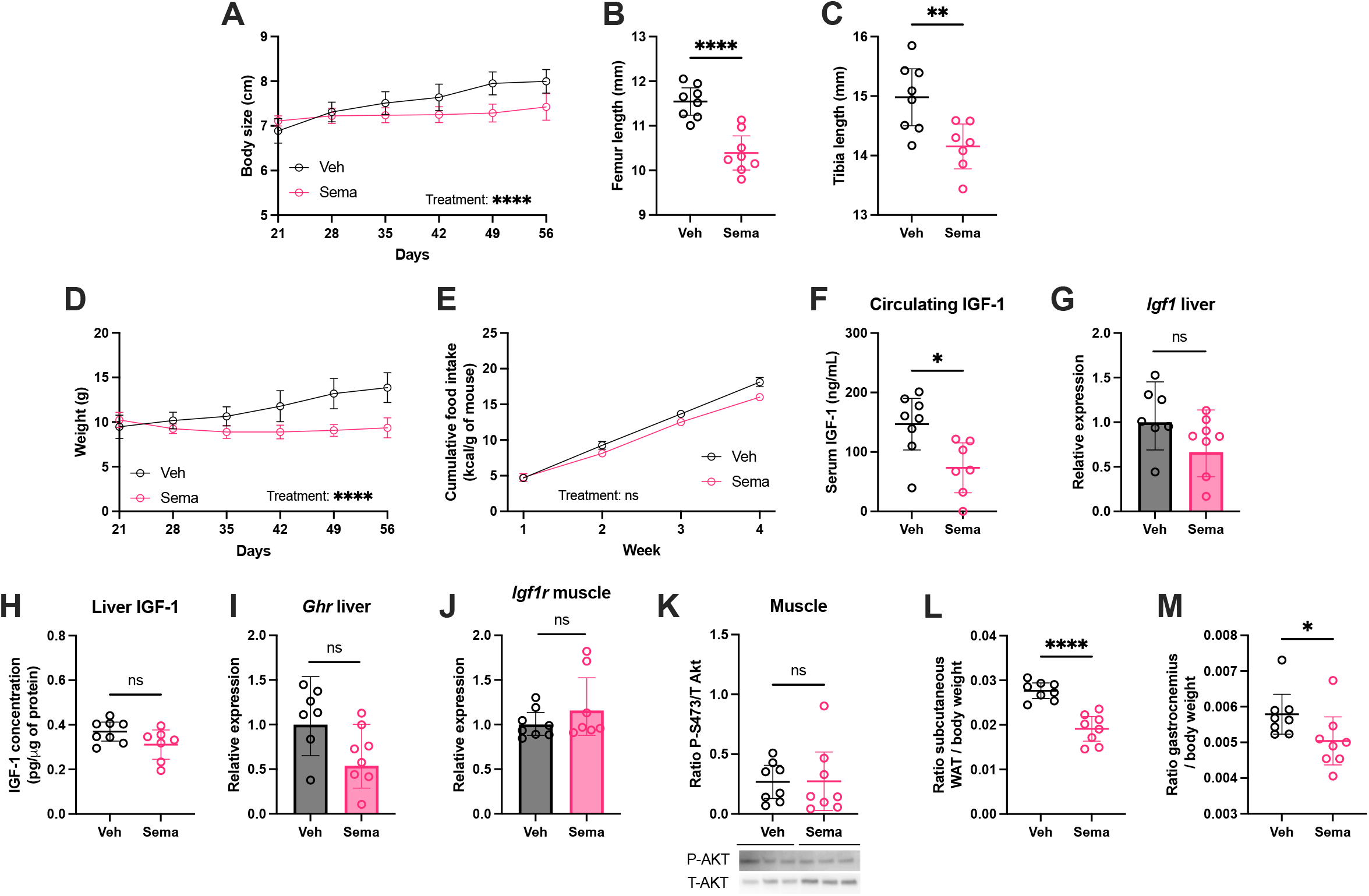
Semaglutide further impairs linear growth and alters body composition in stunted juvenile mice. (A) Body size. Femur (B) and tibia (C) lengths at postnatal day 56 (P56). (D) Body weight over time. (E) Cumulative food intake. (F-K) Somatotropic axis markers: serum IGF-1 concentration (F), liver *Igf1* expression (G), liver IGF-1 concentration (H), liver *Ghr* expression (I), muscle *Igf1r* expression (J) and phosphorylated Akt (S473) to total Akt ratio (K) measured by Western blot and normalized to total protein content. (L-M) Relative weights of subcutaneous WAT (L) and gastrocnemius muscle (M) at P56. Veh, vehicle-treated; Sema, semaglutide-treated. Data are shown as mean ± 95% CI, except for cumulative food intake (mean ± SEM). *p < 0.05, **p < 0.01, ****p < 0.0001, ns: not significant.

Semaglutide is known to reduce body weight, largely via appetite suppression, in patients with obesity^5,6^ and in adult mouse models^30,31^. In SID-fed juveniles, semaglutide reduced weight gain (32% lower than vehicle-treated counterparts), but notably did not reduce food intake (Fig. 2D-E), indicating that the observed growth impairment is not attributable to reduced caloric intake.

Given previous reports of GLP-1 receptor agonists interfering with IGF-1 signaling^16^, we examined the somatotropic axis in SID-fed mice treated with semaglutide. Circulating IGF-1 levels were reduced in semaglutide-treated animals (Fig. 2F), consistent with impaired growth. However, liver expression of *Ghr* and *Igf1*, as well as hepatic IGF-1 protein levels, were unchanged between semaglutide- and vehicle-treated animals (Fig. 2G–I), arguing against a disruption of GH signaling at the hepatic level. Since IGF-1 promotes anabolic processes in peripheral tissues such as muscle^32^, we further assessed IGF-1 signaling in this tissue. Semaglutide treatment did not affect *Igf1r* expression or downstream AKT phosphorylation in skeletal muscle (Fig. 2J-K). Taken together, these results do not support major alterations in somatotropic axis function as the primary mechanism underlying semaglutide’s growth-inhibitory effects in stunted juveniles.

Interestingly, semaglutide-treated SID-fed mice displayed altered body composition, with lower relative weights of subcutaneous white adipose tissue and gastrocnemius muscle (Fig. 2L-M). In addition, semaglutide improved glycemic response to OGTT and lowered fasting glycemia, similar to effects observed in CD-fed animals (Fig. S2A-B). However, fasting insulinemia and HOMA-IR were not significantly altered (Fig. S2C-D). Taken together, these findings demonstrate that chronic semaglutide treatment modulates energy metabolism and body composition in growth-impaired juveniles, independently of food intake or major changes in somatotropic axis activity.

### Semaglutide alters energy metabolism–related gene expression in metabolically active tissues of stunted juvenile mice

To better understand how semaglutide impacts growth-compromised juveniles, we performed transcriptomic analysis of the liver, a central organ in both metabolic and growth regulation. Among ~13,000 detected genes, only 126 were differentially expressed between semaglutide- and vehicle-treated animals (47 upregulated, 79 downregulated) with no signature of alterations in metabolic-or growth-related processes, indicating relatively mild hepatic transcriptional reprogramming (Fig. 3A).

**Figure 3.**
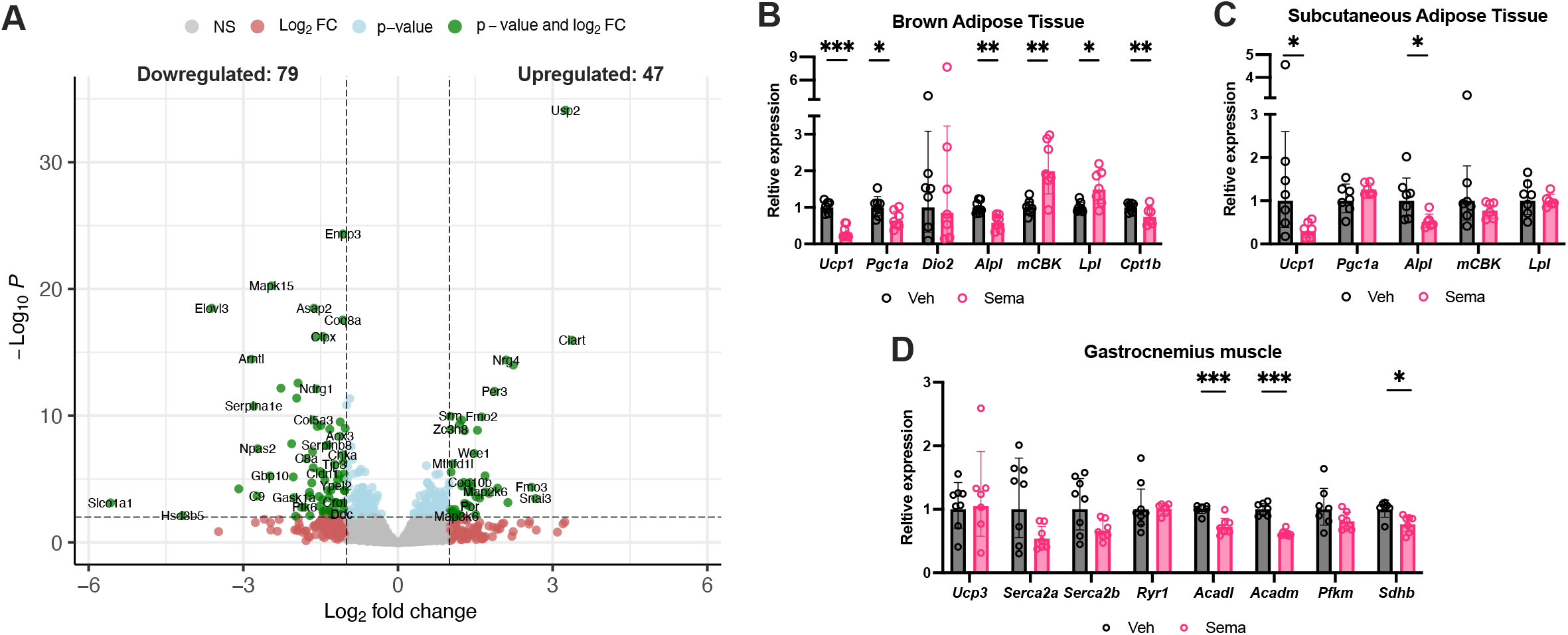
Semaglutide alters energy metabolism gene expression in metabolically active tissues in stunted juvenile mice. (A) Volcano plot showing differential gene expression in the liver of SID-fed juveniles treated with semaglutide (N = 5) versus vehicle (N = 4). (B-D) Relative expression of energy metabolism-related genes in brown adipose tissue (BAT, B), subcutaneous WAT (C), and gastrocnemius muscle (D). The reference genes used to normalize gene expression are: *Rpl32* and *Rsp18* for BAT and WAT; *Gapdh* and *Tbp* for muscle. Veh, vehicle-treated; Sema, semaglutide-treated. Data are shown as mean ± 95% CI. *p < 0.05, **p < 0.01, ****p < 0.0001.

Given the limited hepatic response, we next examined semaglutide’s impact on selected gene’s expression profiles in peripheral metabolically active tissues that are more direct targets of GLP-1 signaling: brown adipose tissue (BAT), subcutaneous white adipose tissue (scWAT), and gastrocnemius muscle. We focused on genes involved in key energy-expending pathways, including fatty acid oxidation (*Cpt1b, Lpl, Acadm, Acadl*), thermogenesis (*Ucp1, Ucp3, Pgc1a, Dio2*), basal energy metabolism (*Sdhb, Pfkm*), and energy-wasting futile cycles (*Alpl, mCBK, Serca2a, Serca2b*).

In BAT (Fig. 3B), semaglutide-treated mice showed reduction of thermogenesis-related genes (*Ucp1* and *Pgc1a*) and *Cpt1b*, suggesting decreased thermogenic and oxidative activity. Interestingly, *Lpl* expression was slightly increased, possibly reflecting a compensatory mechanism—despite reduced thermogenic drive. Expression of futile cycle markers was mixed, with decreased *Alpl* and increased *mCBK*, suggesting altered energy-wasting activity, though the direction of change remains unclear. These data point to altered energy expenditure in BAT following semaglutide treatment in stunted juveniles.

In scWAT (Fig. 3C), *Ucp1* was similarly downregulated, indicating reduced beigeing capacity. In gastrocnemius muscle (Fig. 3D), we observed decreased expression of *Acadl* and *Acadm* (fatty acid oxidation enzymes), along with lower *Sdhb* levels, a marker of mitochondrial oxidative capacity. Collectively, these transcriptional changes evoke that semaglutide may suppress catabolic and energy-expending pathways along multiple peripheral tissues.

Catabolic activity is essential during growth tissues to sustain the high energy requirements of tissue development. Therefore, reduced thermogenic and oxidative capacity in semaglutide-treated stunted juveniles may contribute to an overall energy-conserving phenotype that is incompatible with normal tissue growth.

## Discussion

Understanding the potential impacts of the increasingly prescribed GLP-1 receptor agonist semaglutide in the clinics is essential to guarantee its safety, particularly as growing evidence highlights its broad physiological effects. However, the mechanisms underlying some of these actions remain poorly defined. Given the expanding use of semaglutide in pediatric populations, we aimed to investigate its interaction with a key developmental process: linear growth.

In juvenile mice raised under nutritionally optimal conditions, semaglutide enhanced glucose clearance, consistent with effects described in adult rodents and humans^3,33^. Notably, these metabolic effects were uncoupled from changes in energy balance considerations, as food intake and weight gain remained unchanged. To our knowledge, this is the first study to report a dissociation between semaglutide’s effects on glucose homeostasis and weight gain in juveniles, suggesting age-specific mechanisms of action^17^. Chronic semaglutide treatment did not affect linear growth in these animals, consistent with clinical studies in children with obesity treated with semaglutide or liraglutide, which reported no evidence of altered growth outcomes^7,34,35^.

As linear growth is a key indicator of juvenile health, we explored the potential interaction between GLP-1 receptor agonists and the somatotropic growth axis^16^, in the context of growth disruption. Using a model of diet-induced stunting, we found that semaglutide further impaired linear growth of those juveniles, despite maintaining its beneficial effects on glucose management. However, our investigation of the GH/IGF-1 somatotropic axis did not reveal major alterations in hormone production and signaling, challenging our initial hypothesis that semaglutide’s effects on growth might be mediated through disruption of this pathway.

Semaglutide has been shown to influence many physiological processes regulating energy expenditure^36–38^. Given that semaglutide- and vehicle-treated stunted juveniles had similar food intake, we hypothesized that the observed growth impairment could be due to reduced energy availability, rather than decreased intake. Transcriptomic and gene expression analyses in metabolically active tissues revealed signs of decreased catabolic activity and suppressed thermogenesis, particularly in BAT. These findings suggest that semaglutide treatment in stunted juveniles leads to a shift in energy allocation—reducing energy-expensive processes such as tissue anabolism and thermogenesis, which may impair growth.

In conclusion, our study reveals unexpected consequences of chronic semaglutide treatment during juvenile development in mice, particularly under growth-limiting nutritional conditions. Our findings challenge the notion that GLP-1R agonist–mediated improvements in glucose metabolism are solely driven by reduced food intake and highlight the importance of developmental timing and nutritional context when evaluating safety and efficacy of GLP-1 receptor agonists. These results underscore the need for further studies to investigate the integrated mechanisms of action of GLP-1 receptor agonists across physiological states and life stages.

## Acknowledgments

The authors thank Benjamin Gillet and Sandrine Hughes from the sequencing platform of IGFL (PSI) for their help regarding the sequencing project. They also thank Raghav Galgali for his help in setting up the bioinformatics pipeline for liver transcriptomic data analysis, the animal facility of IGFL (PEHR) where animal experiments were conducted, and the associated group for animal welfare (SBEA).

## Data availability

The RNA sequencing dataset generated in this study has been deposited in the NCBI Gene Expression Omnibus (GEO) under accession number GSE266477.

Some or all datasets generated during and/or analyzed during the current study are not publicly available but are available from the corresponding author on reasonable request.

## Supplementary figure captions

**Figure S1.**
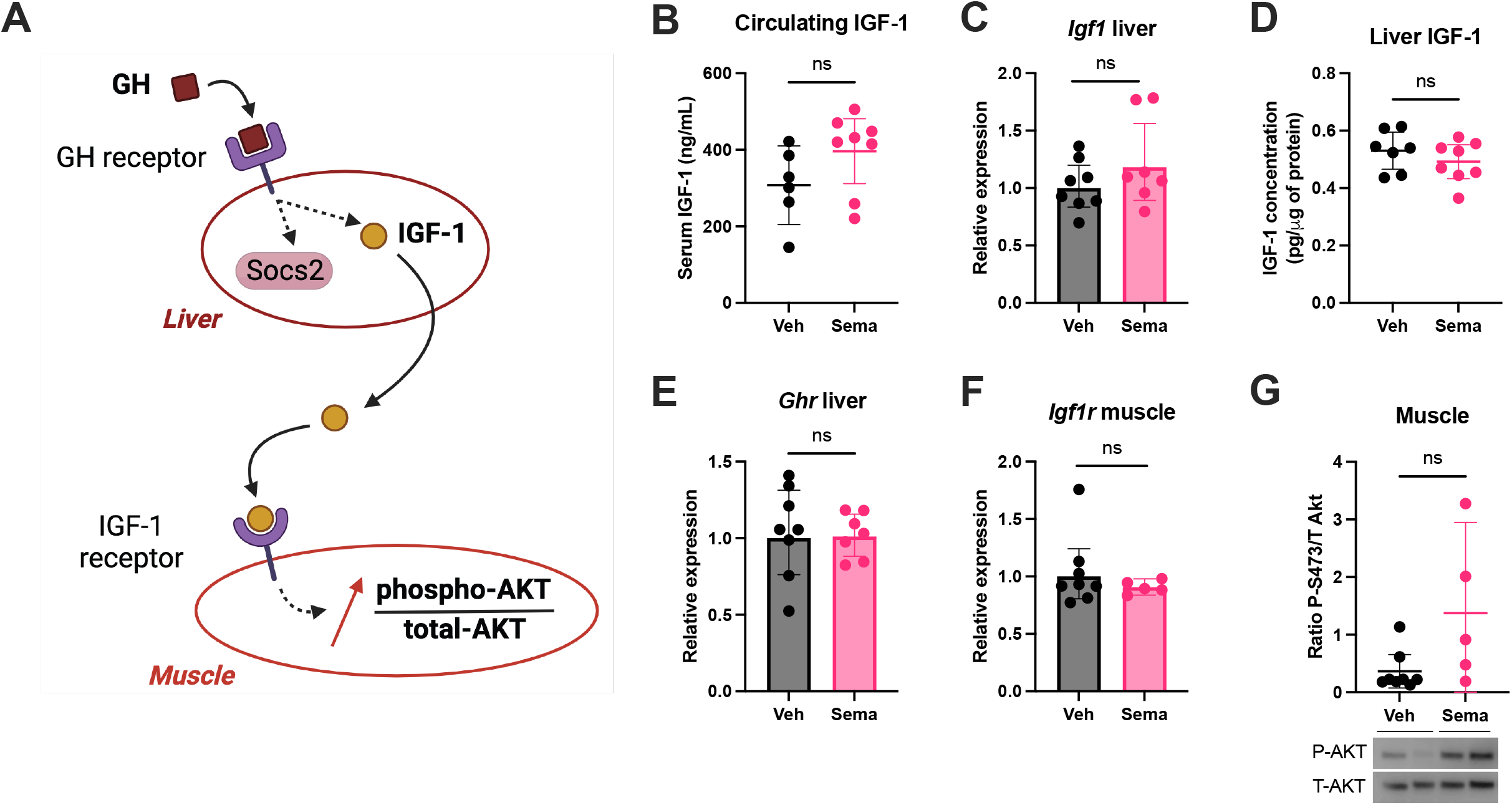
Semaglutide treatment does not alter the somatotropic axis in control-fed juvenile mice. (A) Schematic representation of the somatotropic GH/IGF-1 axis. (B-G) Somatotropic axis markers at P56: serum IGF-1 (B), liver *Igf1* expression (C), liver IGF-1 concentration (D), liver *Ghr* expression (E), muscle *Igf1r* expression (F), and phosphorylated Akt (S473) to total Akt ratio (G) measured by Western blot and normalized to total protein content. Veh, vehicle-treated; Sema, semaglutide-treated. Data are shown as mean ± 95% CI. ns: not significant.

**Figure S2.**
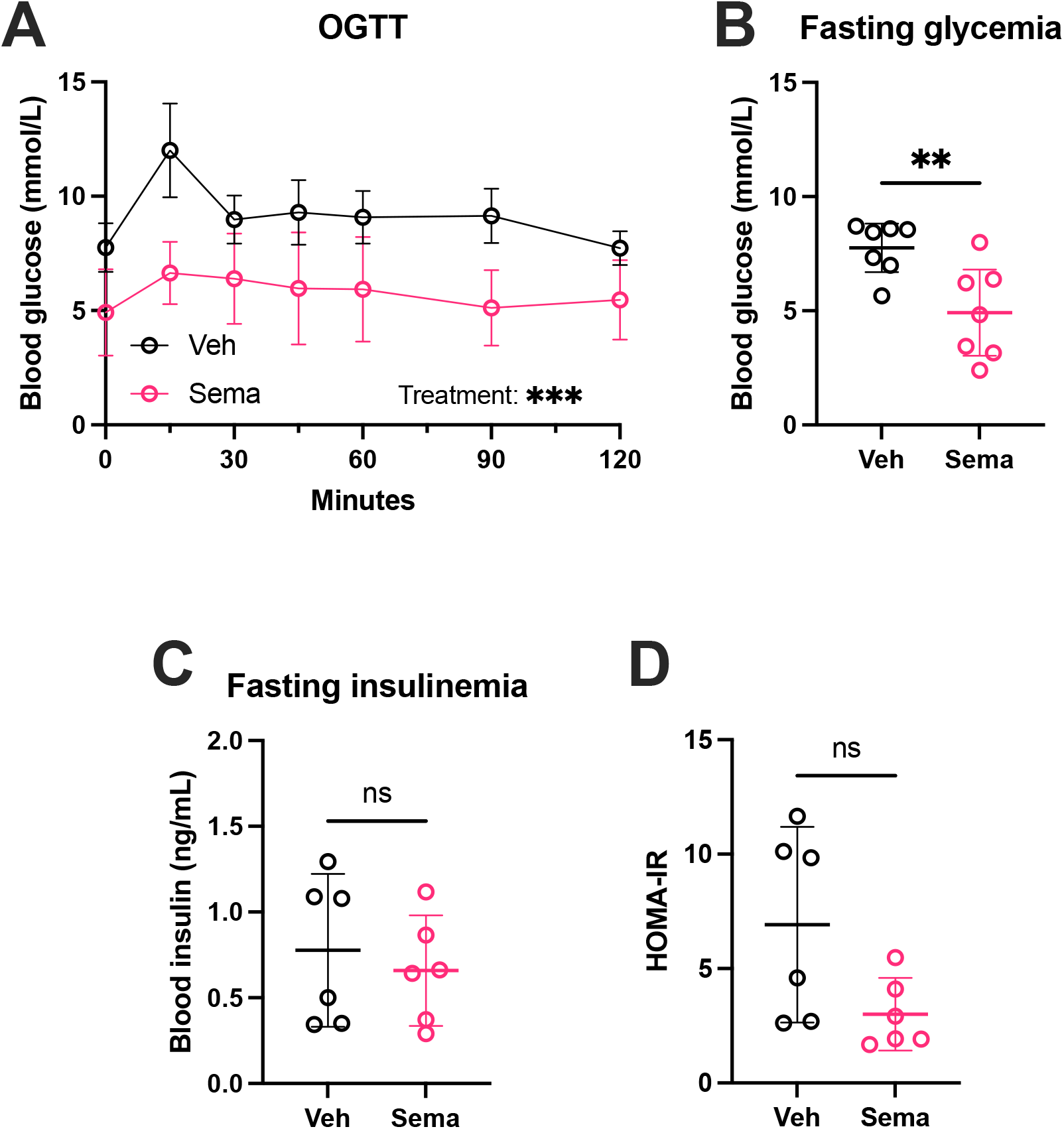
Semaglutide treatment lowers blood glucose and enhances glucose clearance in stunted juvenile mice. (A) Glycemia during an oral glucose tolerance test (OGTT). (B-C) Fasting glycemia (B) and insulinemia (C). (D) HOMA-IR. Veh, vehicle-treated; Sema, semaglutide-treated. Data are shown as mean ± 95% CI. **p < 0.01, ***p < 0.001, ns: not significant.

